# Dissecting human monoclonal antibody responses from mRNA- and protein-based XBB.1.5 COVID-19 monovalent vaccines

**DOI:** 10.1101/2024.07.15.602781

**Authors:** Raianna F. Fantin, Jordan J. Clark, Hallie Cohn, Deepika Jaiswal, Bailey Bozarth, Alesandro Civljak, Vishal Rao, Igor Lobo, Jessica R. Nardulli, Komal Srivastava, Jeremy Yong, Robert Andreata-Santos, Kaitlyn Bushfield, Edward S. Lee, Gagandeep Singh, PVI Study Group, Steven H. Kleinstein, Florian Krammer, Viviana Simon, Goran Bajic, Camila H. Coelho

## Abstract

The emergence of highly contagious and immune-evasive severe acute respiratory syndrome coronavirus 2 (SARS-CoV-2) variants has required reformulation of coronavirus disease 2019 (COVID-19) vaccines to target those new variants specifically. While previous infections and booster vaccinations can enhance variant neutralization, it is unclear whether the monovalent version, administered using either mRNA or protein-based vaccine platforms, can elicit *de novo* B-cell responses specific for Omicron XBB.1.5 variants. Here, we dissected the genetic antibody repertoire of 603 individual plasmablasts derived from five individuals who received a monovalent XBB.1.5 vaccination either with mRNA (Moderna or Pfizer/BioNtech) or adjuvanted protein (Novavax). From these sequences, we expressed 100 human monoclonal antibodies and determined binding, affinity and protective potential against several SARS-CoV-2 variants, including JN.1. We then select two vaccine-induced XBB.1.5 mAbs, M2 and M39. M2 mAb was a *de novo,* antibody, i.e., specific for XBB.1.5 but not ancestral SARS-CoV-2. M39 bound and neutralized both XBB.1.5 and JN.1 strains. Our high-resolution cryo-electron microscopy (EM) structures of M2 and M39 in complex with the XBB.1.5 spike glycoprotein defined the epitopes engaged and revealed the molecular determinants for the mAbs’ specificity. These data show, at the molecular level, that monovalent, variant-specific vaccines can elicit functional antibodies, and shed light on potential functional and genetic differences of mAbs induced by vaccinations with different vaccine platforms.

**GRAPHICAL ABSTRACT:** 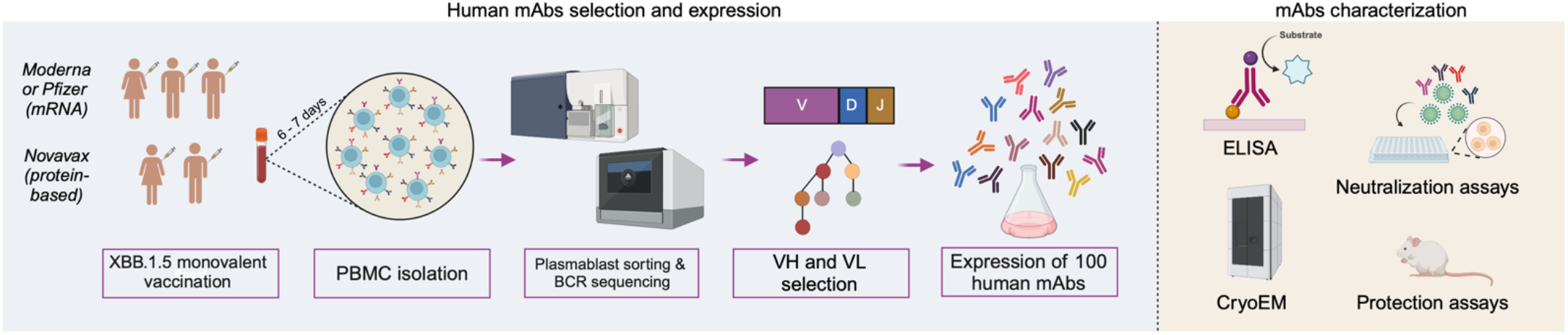

## INTRODUCTION

The spike (S) protein of SARS-CoV-2 has proven to be an essential target for human- neutralizing antibodies(*1–8*). However, spike continues to accumulate mutations, creating new viral variants of concern (VOCs) that often exhibit increased transmissibility and a greater ability to evade the host immune system(*9–11*). Specifically, the Omicron variant of SARS- CoV-2 accrued multiple amino acid mutations within spike and became particularly resistant to neutralization following primary vaccination(*9, 12–14*). Omicron’s evasion of previously elicited antibodies and its development into further antibody-resistant subvariants led to the development of bivalent vaccines(*15*). Among the most recent subvariants, XBB.1.5 became widely recognized for containing key mutations that allow evasion of host immune responses induced by booster vaccinations(*16, 17*). To target the circulating XBB.1.5 strain, the US Food and Drug Administration (FDA) approved updated mRNA monovalent vaccines (*Pfizer or Moderna*) along with a protein-based version adjuvanted with Matrix M^TM^ (*Novavax*) in the fall of 2023(*18, 19*). It is not yet clear whether the functional monoclonal antibodies elicited in response to monovalent Omicron-specific vaccines targeting XBB.1.5 SARS-CoV- 2 represent *de novo* responses or reflect cross-reactivity acquired from previous exposures. Therefore, it became critical to evaluate the ability of monovalent SARS-CoV-2 Omicron- specific vaccines to produce antibodies that recognize those viral variant epitopes. Here, we assessed the human antibody repertoire of plasmablasts from adult participants receiving the mRNA-based (Pfizer, Moderna; 3 participants) or protein-based (Novavax; 2 participants) XBB.1.5 SARS-CoV-2 vaccine. This work reveals novel epitopes of *de novo* and cross-reactive (ancestral and XBB.1.5) antibodies elicited by monovalent XBB.1.5-specific SARS-CoV-2 immunizations.

## RESULTS

### IGHA and IGHG are the predominant isotypes elicited by XBB.1.5 SARS-CoV-2 monovalent vaccines

The antibody repertoire was obtained from five different participants (sex ratio of 1.5:1; male:female). They all had 5-6 prior SARS-CoV-2 vaccinations (primary ancestral scheme consisting of two doses, ancestral SARS-CoV-2, and bivalent booster vaccinations, and the XBB.1.5 monovalent vaccine) (**Fig. 1**). After pairing and filtering 213 mRNA vaccine-induced and 390 protein vaccine-induced paired heavy and light chain sequences of single cell plasmablasts (**Table S1**), we compared somatic hypermutation (SHM) levels in the heavy and light chain V genes between the protein and mRNA groups. For both the heavy and light chains (kappa and lambda), the highest SHM rates were found in the protein-based group **(Fig. S1a;** p ≤ 0.0001). VH mutations in the recombinant protein vaccine group had a median frequency of 8.3%. The mRNA vaccine group presented a lower median mutation frequency of 5.9%, with the top 25% of cells ranging from 8.1% to 15.6%. VL kappa and lambda median mutation frequencies in the recombinant protein vaccine group were also the highest (5% and 5.2%, respectively), with the top 25% of cells ranging between 7.5-25.3% and 7-15.1% respectively. The mRNA vaccine cells showed even lower median mutation frequencies for light chain: 2.6% and 3.8% for VL kappa and lambda, respectively, with the top 25% of cells varying between 4.5-28% and 5.2-10%.

**Figure 1.**
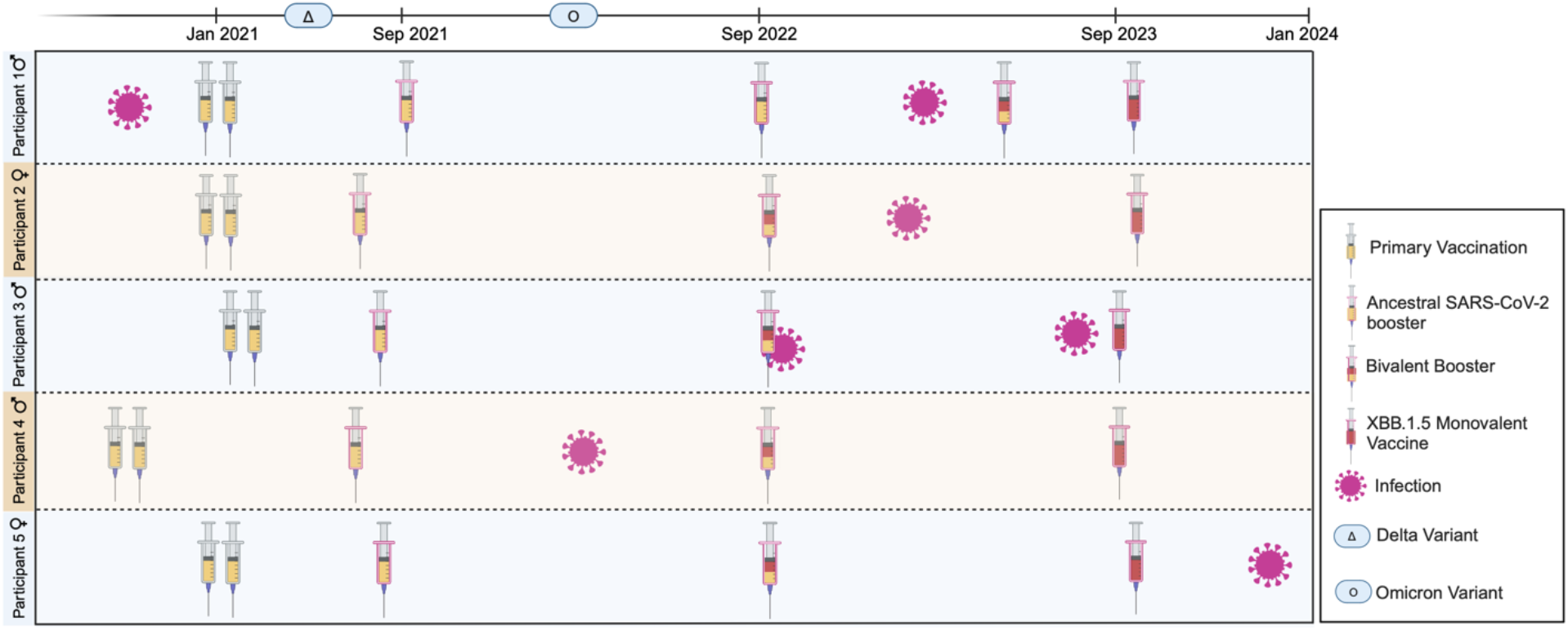
SARS-CoV-2 Immune history of enrolled participants. Participants 1, 2, and 3 received XBB.1.5 mRNA vaccination, while 4 and 5 received the recombinant protein XBB.1.5 vaccine. Delta and Omicron variant’s timeline reflects the United States of America. Created with BioRender.com.

IGHA and IGHG are predominant isotypes in the repertoires of both groups, ranging from 42% to 58.5%; and 26.4% to 54.5% in the mRNA group, respectively. Similarly, in the recombinant protein vaccine-induced plasmablasts, IGHA constitutes 50% - 57.9%, while IGHG constitutes 34.3% - 45.5% of the BCR repertoires **(Fig. S1b).** The highest proportion of IGHM was seen in the mRNA vaccine-induced plasmablasts, ranging from 3.41% to 15.1%. For the protein group, IGHM ranged from 4% to 6.7%.

The highest median mutation frequencies in both groups were found in IGHA cells: 6.9% in the mRNA vaccine-induced plasmablasts and 8.7% in the recombinant protein vaccine-induced plasmablasts **(Fig. S1c)**. At the IGHG1 isotype level, the recombinant protein vaccine-induced plasmablasts also showed the highest SHM frequency values, with significant differences between the groups **(Fig. S1d**; p ≤ 0.001, Mann-Whitney U test).

Clonal groups were determined by clustering BCR sequences that belonged to the same V and J genes and had the same junction lengths, with a Hamming distance threshold of 0.15 applied for the nucleotide sequence of the heavy chain CDR3 regions. In the mRNA vaccine- induced plasmablasts, we identified 164 clonal groups, encompassing 76.9% of the total repertoire sequences, while the recombinant protein vaccine-induced plasmablasts had 289 clonal groups, representing 74.1% of the total repertoire sequences. Expanded clones accounted for 5.2% of BCR sequences in the mRNA vaccine-induced plasmablasts and 6.6% in the recombinant protein vaccine-induced plasmablasts.

Due to the high variability and critical antigen-binding function of the B cell heavy chain CDR3 region, we examined both vaccine groups for physicochemical properties at the amino acid level of the CDR3 region. The recombinant protein vaccine-induced plasmablasts showed a slightly longer mean CDR3 region length than the mRNA vaccine-induced plasmablasts **(Fig. S1e**; p ≤ 0.05, Mann-Whitney U test). Additionally, physicochemical properties such as the aliphatic index **(Fig. S1f**; p ≤ 0.05, Mann-Whitney U test) were significantly altered, with a slightly higher mean in the recombinant protein vaccine induced plasmablasts. Conversely, the mRNA vaccine-induced plasmablasts exhibited heavy chain CDR3 regions substantially more enriched in aromatic amino acids **(Fig.S1g**; p ≤ 0.01, Mann- Whitney U test).

In summary, these results indicate that plasmablasts from both vaccine groups have similar isotype compositions, with IGHA and IGHG being predominant.

#### Assessment of the binding capacity of purified mAbs revealed multiple cross-reactive and one XBB.1.5-specific antibody

Participants were divided into groups based on the XBB.1.5 vaccine type: mRNA (Pfizer or Moderna) or recombinant protein (Novavax), and paired VH-VL sequences were selected if they either: (i) belonged to clonally expanded groups; or (ii) exhibited low levels of somatic hypermutation, enhancing the likelihood of identifying *de novo* antibodies specific for XBB.1.5. Based on these criteria, we synthesized and expressed 100 mAbs (50 from the mRNA vaccine group and 50 from the recombinant protein vaccine group) that were then screened for binding **(Fig. S2)**.

To assess the binding capacity of the mAbs, we tested them in ELISA against ancestral, XBB.1.5, and JN.1 SARS-CoV-2 spike as well as against ancestral, XBB.1.5 and JN.1 RBD. In addition, we assessed binding to ancestral S1, S2, and NTD and an influenza virus H3 hemagglutinin which served as a negative control.

Eighteen binding mAbs were identified in the mRNA vaccine group: 16 bound to ancestral spike (88.8%), 18 to XBB.1.5 spike (100%), 16 to JN.1 spike (88.8%), five to ancestral RBD (27.8.%), four to XBB.1.5 RBD (22.2%) and three to JN.1 RBD (16.7%) **(Fig. 2a – f)**. One of the mRNA mAbs (M2) bound exclusively to XBB.1.5 Spike (5.5%) indicating that this mAb was likely elicited through the most recent SARS-CoV-2 vaccination **(Fig. 2b)**.

**Figure 2.**
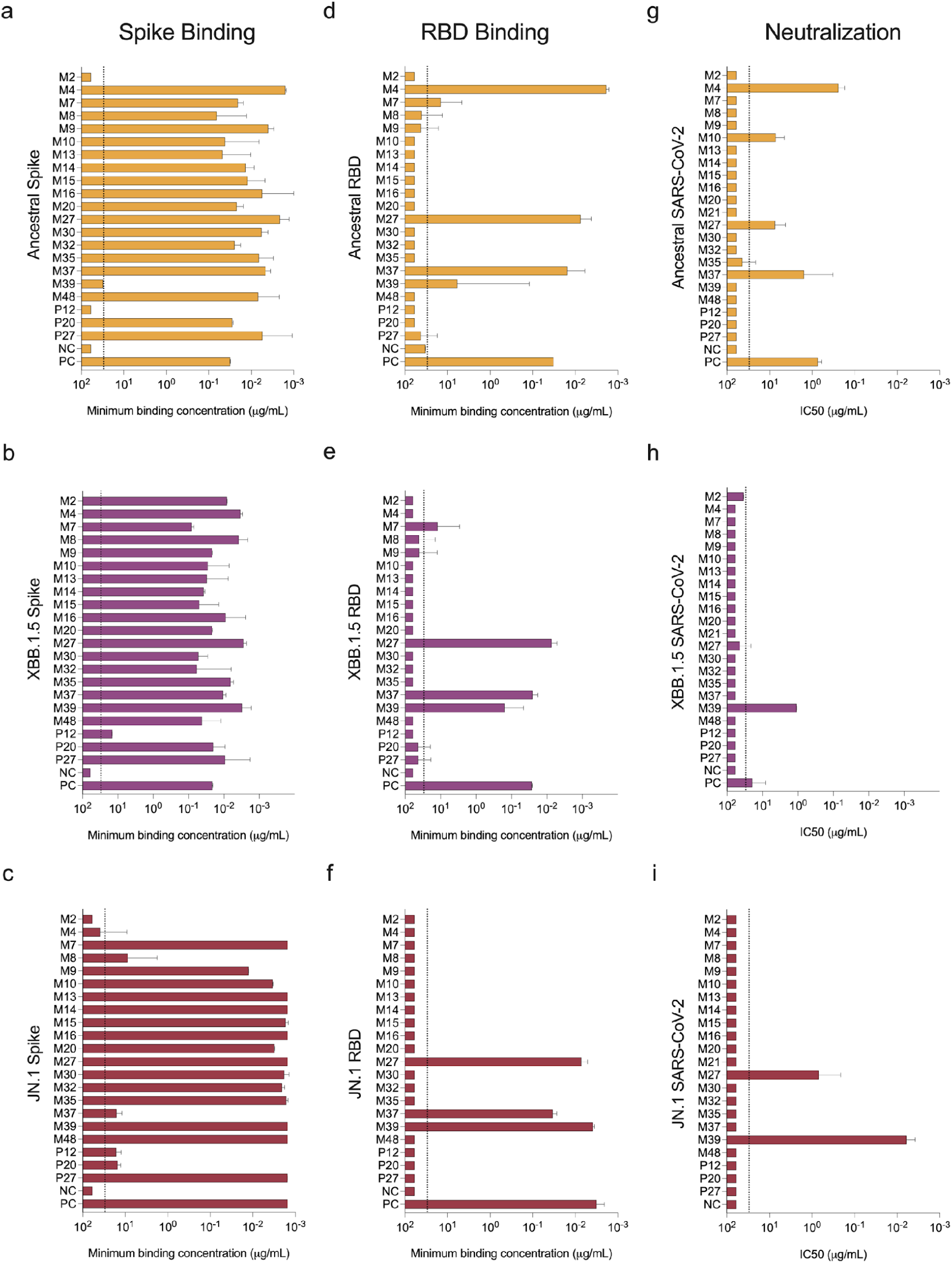
Binding and neutralizing capacity of monoclonal antibodies isolated from participants vaccinated with either mRNA or protein-based SARS-CoV-2 XBB.1.5 vaccines (a – c) Binding activity of the 21 selected mAbs against ancestral, XBB.1.5 and JN.1 spike protein and **(d -f)** ancestral, XBB.1.5 and JN.1 RBD. Neutralizing capacity of binding mAbs against **(g)** ancestral, **(h)** XBB.1.5, and **(k)** JN.1 SARS-CoV-2. Antibodies from the mRNA group are represented by the letter M and protein-based by the letter P. Antibodies that did not bind any antigen are not shown. Only mAbs binding to at least one antigen were selected for the *in vitro* neutralization assays. The dashed line represents the limit of detection (LOD), which is set at the starting dilution of 30 μg/mL. Binding is defined by minimum binding concentration (µg/mL) and neutralization is determined by half-maximal inhibitory concentration (IC50). Negatives were assigned half the LOD.

Three mAbs from the protein group bound to at least one SARS-CoV-2 antigen. Two of these mAbs bound to ancestral spike (66.6%), and all three of them to XBB.1.5 and JN.1 spike (100%) **(Fig. 2a – c)**. There was no binding to the RBDs from any of the SARS-CoV-2 variants tested **(Fig. 2d - f).**

When assessing subunits, 11 (61.1%), 12 (66.6%), and 9 (50%) mRNA vaccine group mAbs bound to ancestral S1, S2, and NTD respectively. Within the recombinant protein vaccinated group, one (33.3%) is bound to S1, and three of them (60%) bound to S2 and NTD **(Fig. S3).**

We selected 21 XBB.1.5 spike mAbs (mRNA; n=18 and protein-based; n=3) for neutralization assays using authentic replication competent SARS-CoV-2 isolates (ancestral, XBB.1.5 and JN.1). Four mAbs efficiently neutralized ancestral SARS-CoV-2, and one out of these four (M27) also neutralized JN.1, but not XBB.1.5. M39 robustly neutralized XBB.1.5 and JN.1, but not ancestral SARS-CoV-2 while M2 did not neutralize any variants *in vitro* (**Fig. 2g – i).** Those three mAbs were then selected for *in vivo* protection assays. M2 as it was the only XBB.1.5-specific mAb (*de novo*); M27 for its broad binding activity and its ability to neutralize both the ancestral strain and JN.1 *in vitro* and M39 for neutralizing both XBB.1.5 and JN.1.

#### M2, M27 and M39 fully protect against XBB.1.5 lethal SARS-CoV-2 challenge in a murine model

We investigated M2, M27, and M39’s ability to protect *in vivo* using the hACE2-k18 lethal SARS-CoV-2 murine challenge model (*20*). Protection was assessed in a prophylactic setting against ancestral SARS-CoV-2, XBB.1.5, and JN.1 variants via the passive transfer of 100mg/kg of mAb intraperitoneally, prior to intranasal infection with a lethal dose of ancestral, XBB.1.5 or JN.1 SARS-CoV-2. M2 (the only XBB.1.5 specific mAb) was protective against XBB.1.5 (100% survival, **Figures 3b and 3e**), but not against ancestral SARS-CoV-2 (Figures 3a and 3d) or JN.1 (**Figures 3c and 3f**). M27 (broadly binder) was capable of robust protection from mortality against all viruses (100% of survival for ancestral and XBB.1.5 and 80% for JN.1) (**Figures 3a – 3f**), despite exhibiting a lack of XBB.1.5 neutralization *in vitro*. For M39 (neutralizing XBB.1.5 and JN.1), full protection was observed when challenged with XBB.1.5, with only partial protection for JN.1 (∼60% of survival) (**Figures 3b – 3f**) and no protection from ancestral SARS-CoV-2 (**Figures 3a and 3d**). Notably, M2 and M27 did not neutralize XBB.1.5 *in vitro* but protected mice following challenge with XBB.1.5 SARS-CoV-2. These data demonstrate that XBB.1.5 vaccine-elicited human mAbs can protect against SARS-CoV-2 challenge in a murine model.

**Figure 3:**
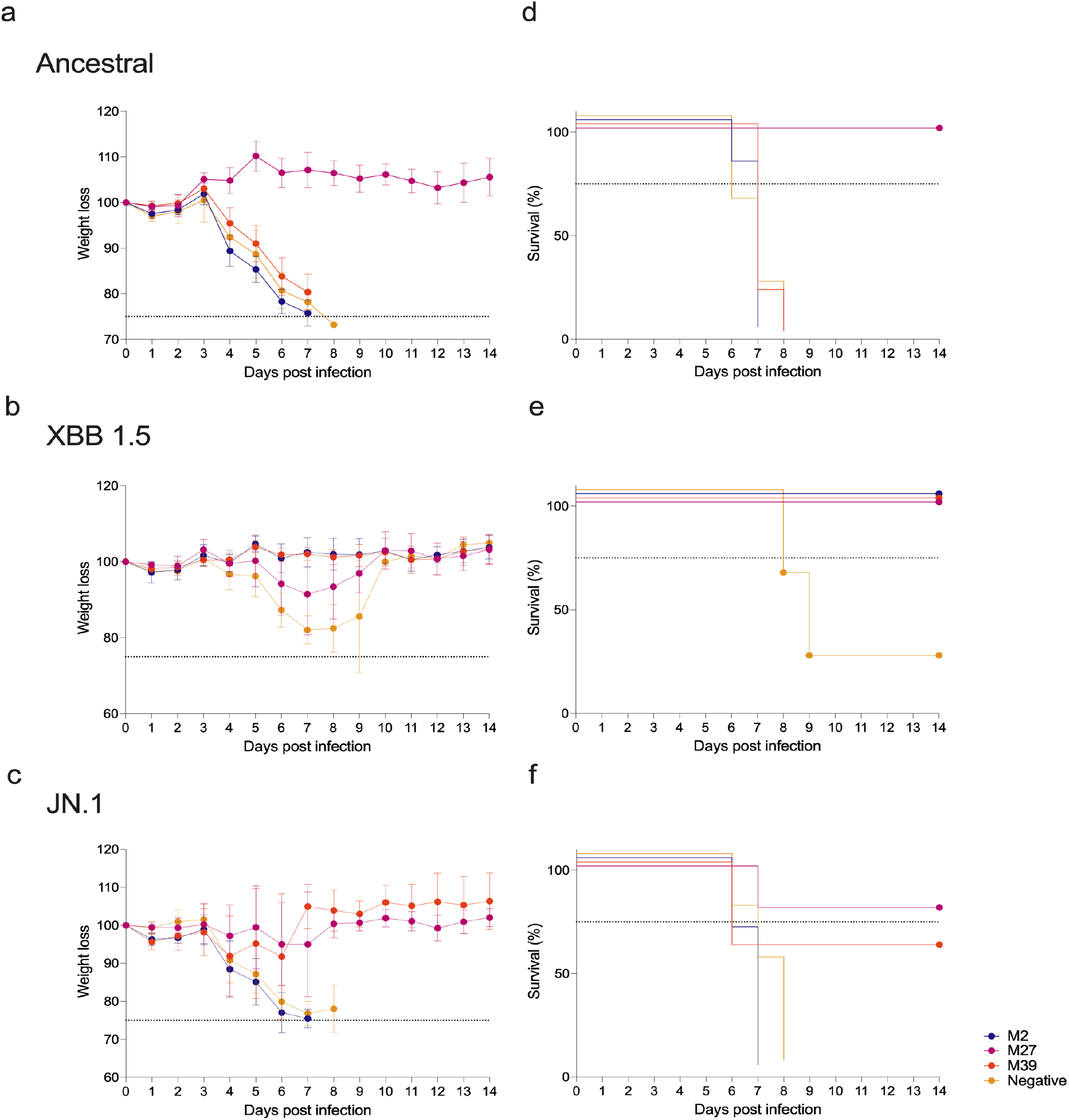
***In vivo* protection by prophylactic treatment with mAbs M2, M27 and M39. (a – c)** Weight loss in mice treated with 10 mg/kg (intraperitoneally) of M2, M27, or M39 mAbs two hours before challenge with a 3xLD50 dose of **(a)** WA1/2020, **(b)** XBB.1.5 and **(c)** JN.1. **(d – f)** Survival curves of mice treated with M2, M27, or M39 mAbs and infected with **(d)** WA1/2020, **(e)** XBB.1.5, and **(c)** JN.1. An influenza virus anti-hemagglutinin mAb, CR9114, was utilized as a negative control. Antibodies from the mRNA group are represented by the letter M and protein-based by the letter P.

#### Cryo-EM structure of M2 Fab and M39 Fab with SARS-CoV-2 XBB.1.5 spike

To characterize the epitope of mAbs M2 and M39, and the intermolecular interactions at the antibody-antigen interface, we used single-particle cryo-EM to determine the structure of M2 Fab and M39 Fab in complex with SARS-CoV-2 XBB.1.5 spike. The structures show that M2 Fab binds to the NTD while M39 binds to the RBD region of spike **(Fig. 4a)**.

**Figure 4.**
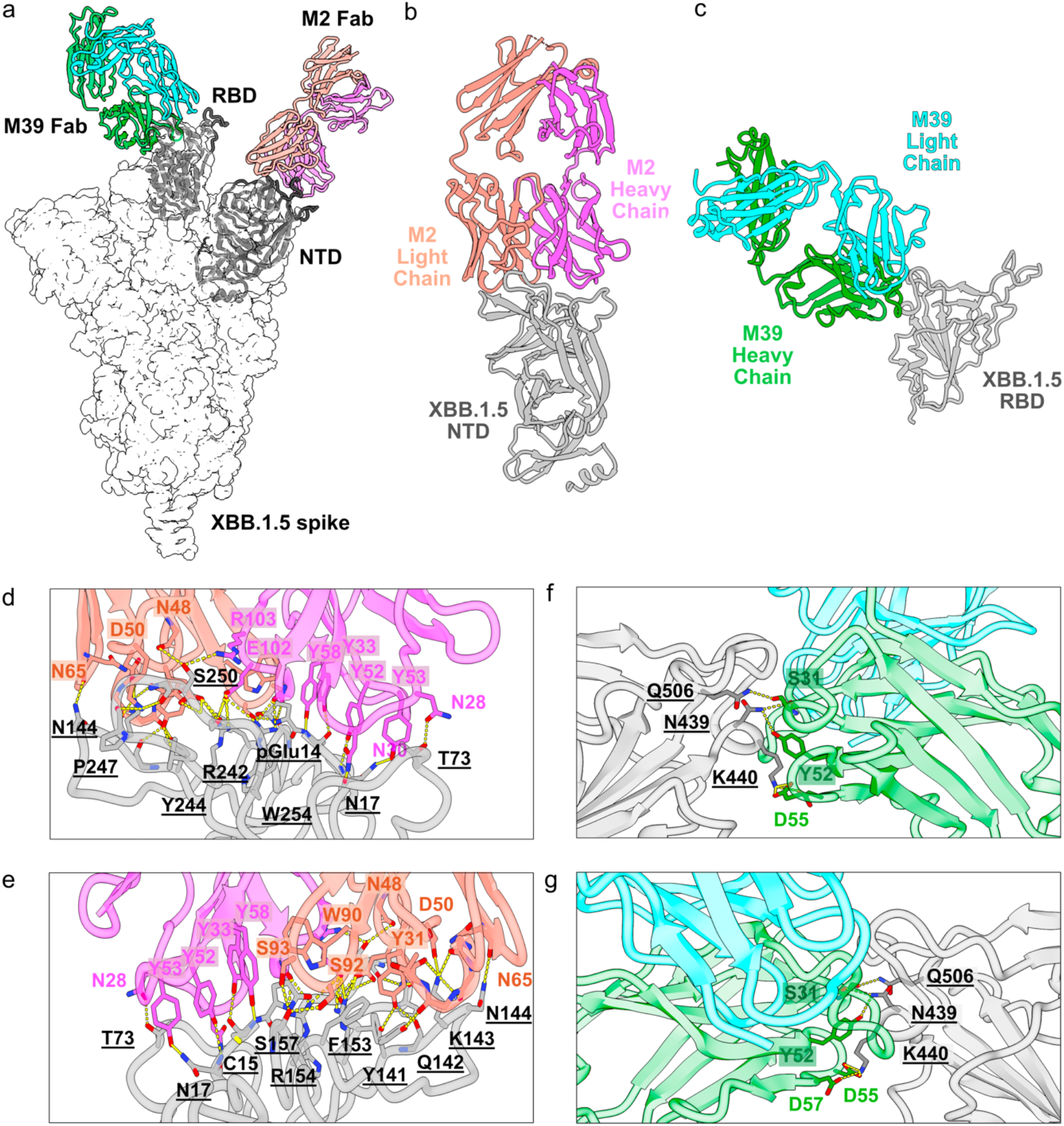
**Cryo-EM structures of M2 and M39 Fabs in complex with XBB.1.5 spike**. **(a)** Surface regions of the SARS-CoV-2 spike contacted by the two antibodies M2 and M39. The global composite cryo-EM map is shown as a transparent surface, with the NTD:M2 Fab and RBD:M39 Fab complexes docked and shown in cartoon representation. **(b)** The M2 Fab defines an epitope on the “top” side of the NTD. M2 creates an extensive surface contact area and engages NTD with both its heavy and light chains. **(d)** and **(e)** are two 180° views along the y-axis that show details of the intermolecular interactions between the M2 Fab and NTD with numerous polar and non-polar interactions. p-GLU designates pyroglutamate. **(c)** The M39 Fab defines an epitope on RBD engaged by numerous published “class 3” antibodies. M39 **(f)** and **(g)** are two 180° views along the y-axis that show details of the M39 Fab:RBD molecular interface with very few interacting residues forming salt bridges.

The global structure of the M2:spike complex shows three Fabs binding to one spike trimer (**Fig. S4**), parallel to its 3-fold axis, and therefore suggests that M2 neutralizes the virus via steric hindrance of the angiotensin-converting enzyme 2 (ACE2) receptor binding. M2 mAb harbors a significant number of somatic hypermutations, with 14 and 12 amino acid mutations in the heavy and light chain V-regions relative to their germline sequences, respectively. Our focus refined map at 2.37 Å nominal resolution permitted us to build an atomic model **(Fig. 4b)** and revealed that M2 engages NTD with both its heavy and light chains. We note a cluster of tyrosine residues, Y33, Y52, Y53, and Y58, encoded by CDRH1 for the first and CDRH2 for the latter three. The four tyrosines engage in extensive polar and non-polar interactions, primarily with the very N-terminus of spike (residues 14-17), including the terminal pyroglutamate (**Fig. 4d-e**). Other notable heavy chain contacts, which arose through somatic hypermutation, include S28N and S30N in the CDR1 **(Fig. 4d-e, S6a).** The antibody light chain contacts NTD with residues derived both through somatic hypermutation and from the germline, including Y48N and D50 **(Fig. 4d, 4e).** An important interacting region on NTD appears to be the 140s loop bound by the M2 light chain CDRL1, including main chain carbonyls from residues 29 and 30 and germline encoded Y31 side chain, as well as CDRL2 including E49 derived through SHM.

The global structure of M39 indicated that it engages the RBD in the “up” as well as “down” configuration. The global cryo-EM map showed three M39 Fabs bound with one spike trimer - with 2 RBDs in the down, and 1 in the up configuration. We used the M39-bound down RBD for further processing **(Fig. S5e)** and conducted local refinement of the M39 Fab/RBD complex to identify the amino acid side-chain contacts at the antibody/antigen interface. The locally refined map, at 2.65 Å nominal resolution, indicated that the M39 has a small footprint on RBD involving CDRH1 and CDRH2 **(Fig. 4c)**. All the polar contacts within the antibody-antigen interface involve the Fab heavy chain, which forms hydrogen bonds with RBD residues N439, K440, and Q506 **(Fig.4f, 4g)**. These residues are conserved between all variants of SARS-CoV-2 except the ancestral WA.1 which has an asparagine substitution at position 440 **(Fig. S6b)**. The primary amine of the K440 side-chain acts as a hydrogen bond donor for D55 and D57 on M39 heavy chain and is thus the major interacting residue, which explains the reduced binding of this antibody with the WA.1 variant. The relatively small binding interface that is accessible on both the up and down RBD conformations likely contributes to the cross-reactivity and potent neutralization activity of M39 antibody against XBB.1.5 and JN.1 SARS-CoV-2 variants.

Collectively, our structural data suggest that somatic hypermutation in both contact and non-contact residues of M2 may have contributed to the development of affinity specifically towards the NTD of XBB.1.5, while the germline sequence likely endowed M39 with a broad binding breadth.

#### V gene pairing analysis reveals that the IGHV4-39/IGLV3-1 pair is specific to XBB.1.5

To further investigate the connection between mAb composition and their binding and functional results, we performed a V gene pairing analysis of our specific and cross-reactive mAbs. We analyzed the heavy and light chain genes and found a prevalent usage of IGHV1- 69 and IGKV3-20. Although these genes did not pair with each other for the binding mAbs, they formed other pairings capable of binding to all subunits and domains in the ancestral S protein **(Figure 5a)** as well as all variants and their respective RBD domains **(Figure 5b)**.

**Figure 5.**
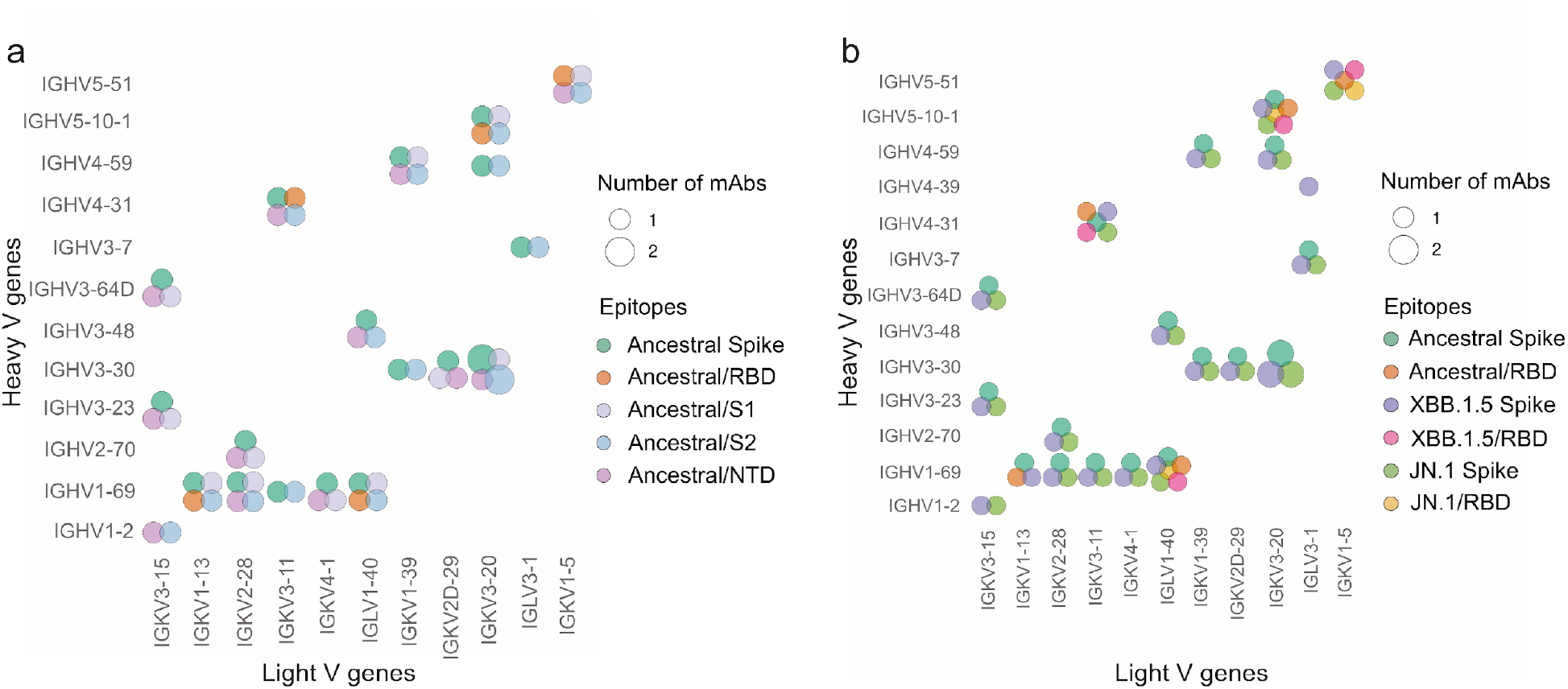
Usage of antibody V gene in heavy and light chain of monoclonal antibodies targeting SARS- CoV-2 spike protein epitopes. (a) V gene pairings for monoclonal antibodies targeting the entire ancestral spike protein, as well as the RBD, NTD, and S1 and S2 subunits. **(b)** V gene pairings for monoclonal antibodies targeting the entire ancestral spike protein, XBB.1.5, and JN.1 variants, along with their respective RBD domains. The size of each circle indicates the number of monoclonal antibodies that use the corresponding V gene pair and bind to the specified epitope. The definition of positive binding was determined based on the cutoffs from Figure 2 and Supplementary Figure S3.

Following IGHV1-69, the IGHV3-30 gene was also frequently used, consistent with previous studies on COVID-19 vaccination(*21, 22*) and infection(*23, 24*). Additionally, we identified two pairs of heavy and light chain V genes previously reported in the literature. The IGHV1- 69/IGKV3-11 pair was used by a public clone that binds to S2, providing partial *in vivo* protection(*25*), and IGHV1-69/IGKV2-28 was previously identified as an RBD binder(*26*), though in our data, the mAb using this pair bound only to the full spike protein in ELISA experiments **(Fig. 5b)**. Finally, we found that the cross-reactive M39 shared the IGHV5- 51/IGKV1-5 V gene combination with an antibody previously identified in naive repertoires of SARS-CoV-2 infected patients (*27*).

## DISCUSSION

Vaccination is the primary and most potent control strategy to prevent COVID-19 effectively and has been proven to elicit robust and protective humoral immunity in humans (*28, 29*). However, SARS-CoV-2 continues to mutate rapidly, developing a wide range of variants capable of immune escape and, thus, causing breakthrough infections(*30*). As of January 2024, CDC reported that the JN.1 lineage had become predominant (∼69% of cases)(*31*), with its subvariants KP.2, KP.3, and LB.1 now dominating different parts of the world. The constant emergence of VOCs demands the steady development and administration of updated vaccines; however, whether these vaccinations can generate functional and variant-specific humoral responses remains unclear.

In our study, most of the antibodies from the mRNA group bound to ancestral spike, XBB.1.5 spike, and JN.1 spike, with only one of them being XBB.1.5-specific, suggesting that the participants’ prior antigenic experiences (besides vaccination, all three participants of the mRNA vaccine group have had infections pre-vaccination) might have influenced their capacity to generate variant-specific responses. As for the recombinant protein vaccine group, none of the mAbs tested were XBB.1.5 specific. Another study that used antibody depletion to assess cross-reactive responses against ancestral and Omicron SARS-CoV-2 variants in participants who have received both primary vaccination and two ancestral/BA.5 or XBB.1.5 matched boosters reported that those without a history of infections had little to no variant-specific antibodies, which seems to be the case for our findings at the monoclonal level for one of the participants(*32–35*).

When designing a SARS-CoV-2 variant vaccine, it is essential to determine how previous exposures to novel variants, such as Omicron, can help to overcome immune imprinting induced by the primary vaccination series. A recent study demonstrated that a single SARS-CoV-2 exposure is insufficient to overcome immune imprinting, as the neutralization of viral variants remained significantly lower compared to the neutralization of the ancestral strain following just one previous Omicron infection(*36*). However, neutralizing antibody titers against viral variants significantly increased following two Omicron infections. Nevertheless, previous infections do not provide absolute protection from emerging SARS-CoV-2 variants, as that study has shown a decrease in the neutralization capacity of sera from Omicron— infected individuals against XBB.1.5 pseudovirus containing targeted escape mutations(*36*). In our study, one of the participants from the mRNA vaccine group had two prior Omicron infections (Participant 3), and all of the *in vitro* neutralizing antibodies (5/5, 100%) came from this participant. The participant’s repertoire also includes an antibody that cross-neutralizes XBB.1.5 and JN.1 (M39); however, although the neutralization capacity of M39 against JN.1 is 100 times higher than XBB.1.5 *in vitro* (IC50: 0.006 for JN.1 and IC50:1.119 for XBB.1.5), the difference is not translated into *in vivo* protection, as 100% of the animals were protected against XBB.1.5 and only 60% against JN.1 (survival). Previous studies have reported that mAbs using the same heavy chain V-gene as M39 (IGHV5-51) also either specifically(*37*) or broadly(*37–40*) neutralize Omicron sub-variants, including XBB. Participant 1, who also had two previous infections (ancestral before primary immunization and Omicron), contributed the only XBB.1.5 specific antibody (M2). This antibody’s robust binding activity (minimal binding concentration < 0.01µg/mL) also translated into robust functional capacity: 100% of survival following murine challenge with XBB.1.5, confirming that experiencing infections before immunization can influence antibody levels and performance(*41*). Consistent with the literature, IGHV1-69 and IGHV3-30 were the most frequently used V genes. These genes are commonly found within the repertoires of vaccinated individuals with a history of infection, as was the case for most of our participants (4/5; 80%)(*42, 43*).

Structural epitope mapping allows us to understand the molecular bases for antibody binding breadth and potentially to predict the location and nature of amino acid substitutions in future viral variants that would facilitate viral immune escape. We defined and analyzed the epitopes of two protective antibodies, M2 and M39, which bind distinct domains on the viral spike – the former binds NTD and the latter RBD. Both mAbs were likely elicited by the XBB.1.5 vaccination or recent infection since they bind the XBB.1.5 variant but do not interact with the ancestral WA.1 spike even though the ELISA data demonstrated some binding. M39 additionally possesses prospective breadth toward JN.1. We posit that insertion 141a (alignment relative to XBB.1.5 sequence) in WA.1 is the main determinant for M2 mAb failing to bind WA.1, and that extensive substitutions in JN.1, including Q142H, F152S and R153G restrict M2 binding to the XBB.1.5 variant **(Fig. S6a)**. Analysis of the M39 structure shows that it binds to RBD outside of the ACE2 binding site and independently of the RBD up or down position, similar to other class 3 antibodies **(Fig. S8)** like S309 and C135(*44, 45*). Although all three antibodies engage a similar epitope involving the residues 439-442 alpha-helix, the interacting residues of the three mAbs on RBD are distinct. M39 and C135 share a relatively small interface area of 562 Å^2^ and 524 Å^2^, respectively, while S309 occupies a larger area of 795 Å^2^. The RBD K440 is the major residue interacting with M39 which might explain the reduced binding of this antibody to the WA.1 variant contains N440. The conservation of the epitope on XBB.1.5, JN.1 and other Omicron variants, however, confers broad cross-reactivity of M39.

Across the assessed metrics, the mRNA vaccine-elicited antibodies were present in a higher proportion and, thus, had a higher number of functionally efficient monoclonals. Overall, the XBB.1.5 vaccine from either platform could not overcome immune imprinting given that most of the antibodies were also ancestral-spike reactive.

Our study shows, at the molecular level, that monovalent, variant-specific vaccines can elicit functional antibodies. It also demonstrates the need for studies that compare the functional and genetic differences of human mAbs induced by immunizations with different vaccine platforms.

### Limitations of the study

Our study has limitations. Although we analyzed over 600 single cells and expressed 100 human mAbs, they spanned five participants. Therefore, we do not have the power to compare individuals vaccinated with mRNA vaccines versus recombinant protein-based vaccines. In addition, we found three antibodies that bound both to S2 and NTD, and we have not been able to elucidate the molecular mechanism of this dual binding so far.

## MATERIAL AND METHODS

### Research Participant Demographic and biospecimen

Biospecimen used in this paper are sourced from two Institutional Review Board (IRB) approved observational research study protocols [STUDY-16-01215/IRB-16-00971 and STUDY-20-00442 /IRB-20-03374] that collect samples before and after viral antigen exposure. Following collection, all specimens were coded for deidentification purposes. Peripheral blood mononuclear cells (PBMCs) were isolated using density gradient centrifugation of whole blood samples from participants and SepMate tubes (STEMCELL Technologies, 85460). PBMCs isolated from the blood samples were stored in liquid nitrogen until analysis. All study participants provided informed consent for participation in research prior to data or sample collection. All human participants’ research is reviewed and approved by the Program for the Protection of Human Participants at the Icahn School of Medicine at Mount Sinai. All human participants research is done in compliance with 45 CFR 46 DHHS regulations.

Biospecimen were collected from five participants following vaccination with mRNA (Pfizer or Moderna) or protein-based (Novavax) vaccine. Sample collection was done between 6-7 days post XBB.1.5 vaccination and cryopreserved peripheral mononuclear cells were used to sort for plasmablasts. The average age of the participant was 46 years old; 3/5 participants were male and all participants ancestries were reported as White. For 4/5 participants, the XBB.1.5 vaccine was the fifth SARS-CoV-2 vaccine dose. XBB.1.5 vaccination was the 6th dose for one participant. 2/5 participants had a homogenous Pfizer mRNA vaccine series. All participants had Pfizer as their primary vaccine type and all participants received at least one bivalent vaccine booster dose. One participant had a SARS-CoV-2 infection prior to any vaccinations. In total, the range of SARS-CoV-2 infections experienced by participants prior to XBB.1.5 vaccination ranged from no infections to two infections.

### Plasmablast sorting

PBMCs were thawed with a combination of warmed Roswell Park Memorial Institute (RPMI) 1640 medium w/ L-glutamine and 25 mM 2-(4-(2-hydroxyethyl)-1-piperazinyl)-ethanesulfonic acid (HEPES) (Corning, catalog no. 10-041-CV), 10% fetal bovine serum (FBS) (Gemini Bio, catalog no.100-106), and 500 units of Benzonase Nuclease HC (Millipore Sigma, 70664-3). After thawing, PBMC were washed with FACS buffer (phosphate-buffered saline (PBS) (Corning, 21-040-CV)) with 2% FBS), resuspended in PBS for counting and viability assessment **(Table S2),** then added to 15mL conical tubes to prepare for staining. Cells were stained with surface antibodies **(Table S3)** diluted in Brilliant Buffer (BD Biosciences, 566349) for 30 minutes at 4 °C protected from light. Following incubation, cells were washed with PBS and dead cells were stained using LIVE/DEAD Fixable Blue Stain Kit (ThermoFisher,

L23105) diluted 1:1000 in PBS and incubated for 15 minutes at room temperature protected from light. Subsequently, cells were washed with FACS buffer and resuspended for sorting using the AuroraCS. Analyses were performed using FlowJo V.10.9 and the gating strategy used to sort plasmablasts is shown in **Figure S7**.

### Single-cell RNA sequencing library preparation

Following the manufacturer’s protocols, sorted plasmablasts were prepared with the Chromium Next GEM Single Cell 5’ Reagent Kit v2 (10x Genomics, 1000244), combined with barcoded Gel Beads (10x Genomics, 1000267), and partitioned into Gel Beads-in-emulsion (GEMs) using a Chromium X controller and the Chromium Next GEM Chip K Single Cell Kit (10x Genomics, 1000286). Following GEM generation, the GEMs underwent reverse transcription to produce cDNA containing an Illumina TruSeq Read 1 sequence, unique molecular identifier (UMI), template switch oligo (TSO), and 10x barcode. The resulting cDNA was then purified using Dynabeads MyOne SILANE (10x Genomics, 2000048) and amplified with reagents from the Chromium Next GEM Single Cell 5’ Reagent Kit v2 (10x Genomics, 1000244) to facilitate B cell receptor (BCR) amplification using a BCR Amplification Kit (10x Genomics, 1000253). Utilizing a Library Construction Kit (10x Genomics, 1000190), the BCR transcripts subsequently went through fragmentation, end-repair, A-tailing, adaptor ligation, and sample index PCR to generate V(D)J libraries containing standard Illumina primers, P5 and P7, dual indices, i5 and i7, and an Illumina TruSeq Read 2 sequence. Following cDNA amplification, BCR amplification, and library construction, samples were purified using SRPIselect (Beckman Coulter, B23317) and quantified on an Agilent Tape Station 4150. Prior to sequencing, the size (bp) of the V(D)J libraries was quantified on an Agilent TapeStation 4150 using a D5000 ScreenTape and Reagents (Agilent, 5067-5588, 5067-5589), while the concentration (ng/uL) was measured on a Qubit 4 using a Qubit 1X dsDNA High Sensitivity (HS) Assay Kit (Invitrogen, Q33231). The V(D)J libraries were then denatured and diluted using a MiSeq Reagent Kit v3 (150- cycle) (Illumina, MS-102-3001) and sequenced on an lllumina MiSeq. Following sequencing, FASTQ files were converted from raw BCL files using the Illumina Basespace platform and processed for data analysis on the 10x Genomics Cloud.

### Bioinformatic analyses of BCR sequencing

We used the Immcantation framework to analyze V(D)J data outputted from 10x Genomics CellRanger. V, D, and J genes were assigned using IgBlast v1.20.0 followed by parsing the output to the Adaptive Immune Receptor Repertoire (AIRR) Community Rearrangement using Change-O v1.3.0. Cells with non-productive sequences, multiple heavy-chains, or only light chains were removed. Among multiple light chains, the one with the highest number of mapped reads was chosen. Cells with no light chain were also removed. Clonal groups were defined using the hierarchicalClones() function from SCOPer package v.1.3.0(*46*), which clusters BCR sequences based on common IGHV gene annotations, IGHJ gene annotations, and junction length. This function defines groups based on sequence similarity, using the within-group junction Hamming distance by single linkage, previously determined by the distToNearest() function, from the SHazaM package v.1.2.0(*47*), and using a similar threshold for the junction region. The threshold was manually selected as 0.15. Clonal groups were additionally formatted using running formatClones from the Dowser package 2.0 (*48*), and identical sequences were collapsed.

We estimated clonal abundance and quantified B cell clonal diversity using the estimateAbundance and alphaDiversity functions from the Alakazam package v.1.3.0(*47*), respectively. For alphaDiversity, we applied resampling strategies to account for variations in sequencing depth by downsampling each sample to the same number of sequences, and at least 50 B cells per participant and 500 repetitions was reported. This method measures diversity as a smooth function of the parameter “q”, which corresponds to the most common diversity metrics in ecology, namely: species richness (q = 0), the Shannon-Weiner index (q = 1), and the Simpson index (q = 2)(*49*).

Somatic hypermutation was determined by first reconstructing germline V and J sequences using CreateGermline.py from Change-O, with masked D segment on downstream analysis. The total number of mutations (R/S mutations in CDRs or frameworks) was determined for the V gene using observedMutations from SHazaM. V gene usage and amino acid properties for the CDR3 region were calculated using, respectively, countGenes and aminoAcidProperties from the Alakazam package.

### VH/VL selection for mAbs expression

Ig V(D)J and C gene assignments for plasmablast BCR sequences were performed with IgBLAST using the AssignGenes and ParseDb functions from Change-O. These were filtered to include cells with unique VH and VL sequences. Among cells with multiple light chain sequences, the light chain sequence with the highest number of reads was chosen. Clonal expansion analysis was performed on the filtered sequences using the DefineClones function from Change-O. Cells belonging to the same expanded clone had identical V and J gene usages in both heavy and light chains, along with >90% similarity in the heavy chain complementarity determining region 3 (CDR3) nucleotide sequence. VH/VL sequences for mAb expression were selected using the following criteria: unique VH/VL sequences within all the expanded clones for mRNA vaccinees and the most mutated sequences in each of the top six (Participant 4) and top four (Participant 5) clonotypes for the protein group were chosen. This is because the number of sequences was higher for the Protein group than the mRNA group, from which we selected all expanded clones. In addition, expanded clones with at least two identical sequences were also selected for both vaccine groups. The definition of unique sequences used was VH/VL sequences that were not part of an expanded clone and had a VH mutation frequency < 5% (to enrich for de novo B cells) and unique VJ gene usages in each subject (mRNA) and both participants combined (protein- based). We chose this SHM filter to enrich for de novo XBB.1.5 specific mAbs, as they are likely to have lower SHM than cross-reactive mAbs. Three sequences from the mRNA group (M13, 14, and 15) were chosen for mAbs for presenting similarity (identical V, D, and J genes and higher than 70% identity in the CDR3 aa sequence) with antibodies from a COVID-19 antibodies database (CoV-AbDab) (*50*).

### Monoclonal antibody (mAb) production

Sequences from all participants were chosen for mAb expression based on clonal expansion as described before. Variable heavy and light chain domains were reformatted to IgG1, synthesized, and cloned into mammalian expression vector pTwist IgG1, IgL2, or IgK utilizing the Twist Bioscience eCommerce portal. Plasmidial DNA of said vectors were then extracted from E.coli using PureYield^TM^ Plasmid Maxiprep System (Promega, A2393) followed by transfections of HEK 293F and EXPI293 cells were carried out using the lipid-based Thermo Fisher Expi293™ Expression System Kit (ThermoFisher, A14525) and performed according to the manufacturer’s protocol. Briefly, plasmidial DNA, ExpiFectamine and 25, 50, or 100mL cell cultures were incubated at 37°C 8%CO2 for 18-22 hours on a platform shaker at 125rpm. Enhancers 1 and 2 were then added and incubated on a platform shaker at 125rpm at 37°C 8% CO2 for an additional 5-6 days. Following incubation, cells were pelleted by centrifugation, the culture medium was filtered (0.45µM) and mixed with protein A agarose resin (Gold Biotechnology, St. Louis, USA, P-400-50) for 12 hours at 4°C on a rotary shaker. On the next day, the mix was loaded into Poly-Prep chromatography columns (Bio-Rad Laboratories, Inc., 7311550), IgG was eluted with glycine and immediately neutralized with 1M Tris pH 8 and 5M NaCl. Purified IgGs were pooled, concentrated, and stored at 4°C short term or -80°C long term for later use.

### Monoclonal antibody enzyme-linked immunosorbent assays (ELISAs)

Recombinant proteins were coated on Immulon 4 HBX 96-well plates (Thermo Scientific) in 50 μL per well at a concentration of 2 μg/mL overnight at 4°C. The following day, plates were washed three times using PBS containing 0.1% Tween-20 (PBST; Fisher Scientific) and blocked using PBST supplemented with 3% non-fat milk (Life Technologies). Monoclonal antibodies were diluted to 30 μg/mL and then serially diluted 1:3 to a final concentration of 1.52x10^-3^ μg/mL in PBST supplemented with 1% non-fat milk (Life Technologies). Following blocking, mAb dilutions were added to the plates and incubated for 1 hour at room temperature. A total of 16 wells per plate were intentionally left blank to serve as a measure of background. Plates were washed three times with PBST and 100 μL of mouse anti-human IgG antibody conjugated to horseradish peroxidase (Sigma, A0293) diluted 1:3000 in 1% non-fat milk PBST was added to each well. Following 1 hour of incubation at room temperature, plates were again washed three times with PBST and developed via the addition of 100 μL of o-phenylenediamine dihydrochloride (OPD; Sigma-Aldrich). Plates were developed for 10 minutes at room temperature and then the reaction was stopped by adding 50 μL 3 M hydrochloric acid (HCl, Fisher) and plates were read using a Synergy 4 (BioTek) plate reader at an optical density (OD) of 490 nanometers.

### Neutralization assays

Vero E6 cells expressing transmembrane protease serine 2 (TMPRSS2, BPS Biosciences, 78081) were cultured in Dulbecco’s modified Eagle medium (DMEM; Gibco) supplemented with 10% fetal bovine serum (FBS), 1% minimum essential medium (MEM) amino acids solution (Gibco, 11130051), 100 units/mL penicillin and 100 μg/mL streptomycin (Gibco, 15140122), and 3 μg/mL puromycin (InvivoGen, ant-pr), and 100 μg/mL normocin (InvivoGen, ant-nr). Viral stocks were verified via sequencing and enumerated using the 50% tissue culture infectious dose (TCID50) method. Microneutralization assays were carried out as previously described(*51*). Briefly, Vero.E6 TMPRSS2 were seeded in 96-well tissue culture plates (Corning, 3340) at a density of 1x10^4^ cells per well. 24 hours later, mAbs were diluted to a starting concentration of 30 μg/mL in 1x minimal essential medium (MEM, Gibco) supplemented with 2% FBS and then serially diluted 1:3 to a final concentration of 0.041μg/mL. 80 μL of mAb dilution was then mixed with 80 μL of SARS-CoV-2 diluted to 10^4^ TCID50/mL and allowed to incubate at room temperature for 1 hour. After 1 hour, 120 μL of the virus/mAb mixture was used to infect the cells for 1 hour. The inoculum was then removed and replaced with 100 μL 1xMEM 2% FBS and 100 μL of antibody dilution. Cells were incubated at 37°C in a 5% CO2 incubator for 48 hours prior to fixation with 10% paraformaldehyde (Polysciences) for 24 hours. Once fixed, the paraformaldehyde was removed, and the cells were permeabilized via the addition of 100 μL PBS supplemented with 0.1% Triton X-100 (Fisher). Cells were permeabilized at room temperature for 15 minutes, after which the 0.1% Triton X-100 was removed, and the cells were blocked via the addition of 100 μL 3% non-fat milk (Life Technologies) diluted in PBS. Cells were then stained for the presence of SARS-CoV-2 nucleoprotein using the mAb 17C7 as previously described(*51*).

### In vivo studies

Animal studies were conducted following Animal Care and Use Committee (IACUC) guidelines, according to approved protocols reviewed by the Icahn School of Medicine at Mount Sinai Institutional Animal Care and Use Committee (IACUC). The 50% lethal dose (LD50) was calculated for WA1/2020, XBB.1.5, and JN.1 by infecting 6–8-week-old hACE2-K18 mice with serial dilutions of virus ranging from 10^5^ to 5 plaque-forming units (PFU), diluted in sterile PBS. For protection studies, mice were treated intraperitoneally with a 10 mg/kg dose of mAb, diluted in 100 µl sterile PBS. Two hours later, mice were anesthetized with 0.15 mg/kg ketamine and 0.03 mg/kg xylazine diluted in water for injection (WFI, Gibco) and intranasally infected with a 3xLD50 dose of SARS-CoV-2 WA1/2020, XBB.1.5, or JN.1 diluted in 50 µl sterile PBS. Mice were monitored for weight loss for 14 days post-infection and any animals falling below the 25% weight loss cut-off were humanely euthanized. The influenza A virus antibody, CR9114, was utilized as a negative control(*52*).

### Cryo-EM sample preparation and data collection

SARS-CoV-2 XBB.1.5 HexaPro spike(*53*) was incubated with M2 or M39 Fabs, at 2 mg/mL with a 1.5 molar excess of fragment antigen binding (Fab) for 20 minutes at room temperature. Immediately before grid preparation, fluorinated octyl-maltoside was added to the preformed complex at 0.02% wt/vol final concentration. Then, 3 μl aliquots were applied to UltrAuFoil gold R1.2/1.3 grids and subsequently blotted for 6 seconds at blot force 0 at 22°C and 85% humidity, then plunge-frozen in liquid ethane using a Vitrobot Mark IV system (ThermoFisher Scientific). Grids were imaged on a Titan Krios microscope operated at 300 kV, equipped with a 15eV energy filter, and a Gatan K3 Direct Electron Detector. 6,034 and 6,112 movies were collected for M2 and M39 antibodies, respectively, with a total dose of 49.883 e-/ Å2/s. Images were collected at a magnification of 105,000, corresponding to a calibrated pixel size of 0.825 Å/pixel, with a defocus range from -0.6 to -2.5μm.

### Cryo-EM data processing

#### M2 complex

Movies were aligned and dose weighted using patch motion correction implemented in cryoSPARC v4.3.1. Contrast transfer function estimation was done in cryoSPARC v4.3.1(*54*) using Patch CTF, and particles were picked with cryoSPARC’s template picker. The picked particles were extracted with a box size of 1024 pixels, with 4× binning, and subjected to a 2D classification. Selected particles underwent two more rounds of 2D classification. An initial model, with two classes, was generated from 856,516 selected particles. The best class, containing 663,608 particles, was selected for further processing. After one round of non- uniform refinement, without imposed symmetry, the particles were subjected to 3D classification with six classes. Four classes were picked for further processing which contained 481,202 particles in total. The particles were re-extracted with a box size of 1024 pixels, binned 2x, and selected for further rounds of non-uniform refinement with local and global CTF refinement, yielding the final global map at a nominal resolution of 2.04 Å. The protomer with the best Fab volume was subjected to local refinement with a soft mask extended by 6 pixels and padded by 12 pixels encompassing the N-terminal domain (NTD) and Fab. Local map resulted in 2.38 Å resolution. The complex underwent another round of 3D classification from which particles from 3 out of 5 classes were merged and subjected to local refinement, yielding a final local map at 2.37 Å. The two half-maps from the local refinement were used for sharpening in DeepEMhancer(*55*). The reported resolutions are based on the gold-standard Fourier shell correlation of 0.143 criterion.

#### M39 complex

Movies were aligned and dose-weighted using MotionCorr2. Contrast transfer function estimation was done in cryoSPARC v4.3.1 using Patch CTF, and particles were picked with cryoSPARC’s blob picker. The picked particles were extracted with a box size of 512 pixels, with 4× binning, and subjected to a 2D classification. Selected particles were then subjected to a second round of 2D classification. An initial model was generated on the 1,193,025 selected particles with three classes. The best class, containing 801,085 particles, was selected for further processing. After one round of non-uniform refinement, without imposed symmetry, the particles were subjected to 3D classification with ten classes. Of these, the best seven classes, containing 552,337 particles in total, were combined, re-extracted without binning with a box size of 512 pixels, and selected for further rounds of non-uniform refinement with local CTF refinement, yielding the final global map at a nominal resolution of 2.25 Å. The protomer with the two best Fab volumes was subjected to local refinement with a soft mask extended by 6 pixels and padded by 8 pixels encompassing the receptor binding domain (RBD) and Fabs. This yielded the local map at 2.69 Å resolution. A second round of local refinement was performed with a soft mask encompassing the RBD and one best Fab. This yielded the final local map at 2.65 Å resolution. The two half-maps from the local refinement were used for sharpening in DeepEMhancer. The reported resolutions are based on the gold-standard Fourier shell correlation of 0.143 criterion.

### Model building and refinement

The DeepEMhancer sharpened focused maps were used for model building. Models for both M2 and M39 complexes were initially built using ModelAngelo(*56*) and then manually adjusted using COOT(*57*). N-linked glycans were built manually in COOT using the glyco extension and their stereochemistry and fit to the map were validated with Privateer(*58*). The model was then refined in Phenix(*59*) using real-space refinement and validated with MolProbity(*60*) (See **Table S4**). The structural biology software was compiled and distributed by SBGrid(*61*).

## Statistical Analysis

Statistical tests were performed in Prism 10 (GraphPad Software, La Jolla, CA, USA) or R (version 4.2.1, The R Foundation for Statistical Computing). Comparisons between vaccine groups (mRNA vs protein) were performed using unpaired Mann-Whitney’s test, as also indicated in the figure legends. Only significant test results are shown in the figures and the P-values are defined as follows: *p* ≤ 0.05: *; *p* ≤ 0.01: **; *p* ≤ 0.001: ***; *p* ≤ 0.0001: ****. The area under the curve (AUC) and minimal binding concentrations were calculated using the average OD plus 5x standard deviation of the blank wells.

## SUPPLEMENTAL INFORMATION

Fig. S1 – S8

Tables S1 – S4 References 45, 62 – 63.

## ACKNOWLEDGMENTS

This work was supported in part through the computational and data resources and staff expertise provided by Scientific Computing and Data at the Icahn School of Medicine at Mount Sinai and supported by the Clinical and Translational Science Awards (CTSA) grant UL1TR004419 from the National Center for Advancing Translational SciencesWe thank the Flow Cytometry Core Facility and staff at Icahn School of Medicine for their assistance. We also thank the study participants and our laboratory members for the helpful and enriching discussions.

## Author Contributions

CHC, FK and VS conceptualized the study. RFF, JCC, HC, DJ, BB, AC, JY, RAS, JN, KS, and KB conducted the experiments. RFF, JCC, HC, DJ, VR, BB, IL, GB, FK, VS and CHC analyzed the data. GB, FK, VS, and CHC supervised and funded the study. VS, KS, and JN provided access to biospecimen and metadata from study participants. EL and SK supervised bioinformatic analyses. GB, FK, VS, and CHC were responsible for the decision to submit the manuscript for publication. RFF, JJC, HC, DJ, IL, VR, GB and CHC wrote the manuscript. All authors interpreted the data, provided critical input, and revised the manuscript.

## Data Availability

Deidentified data may be made available to investigators whose proposed use of the data has been approved by the Icahn School of Medicine Ethics Review Board upon request to the corresponding authors. Participants’ metadata have been deposited to BioProject accession number PRJNA1134144 in the NCBI BioProject database.The EM maps have been deposited in the Electron Microscopy Data Bank (EMDB) under accession codes EMD-45443 and EMD-45444 for the M39 and M2 complexes, respectively. The accompanying atomic coordinates have been deposited in the Protein Data Bank (PDB) under accession codes 9CCI and 9CCJ for the M39 and M2 complexes, respectively. The aligned micrographs are available on the Electron Microscopy Public Image Archive (EMPIAR) under accession numbers EMPIAR-12167 and EMPIAR-12151.

## SUPPLEMENTARY MATERIAL

### PVI study group

In alphabetical order: Harm van Bakel^1,2^, Yuexing Chen^1,2^, Christian Cognigni^1,2^, Dylan Fitzgerald^1,2^, Charles

R. Gleason^1,2^, Ana Gonzalez-Reiche^1,2^, Morgan van Kesteren^1,2^, Giulio Kleiner^1,2^, Neko Lyttle^1,2^, Jacob D. Mauldin^1,2^, Jacob Mischka^1,2^, Brian C. Monahan^1,2^, Emilia Mia Sordillo^5^, Reima Ramsamooj^1,2^

1. Department of Microbiology, Icahn School of Medicine at Mount Sinai, New York, NY, USA

2. Center for Vaccine Research and Pandemic Preparedness, Icahn School of Medicine at Mount Sinai, New York, NY, USA

3. Department of Pathology, Molecular and Cell-Based Medicine, Icahn School of Medicine at Mount Sinai, New York, NY, USA

4. The Global Health and Emerging Pathogens Institute, Icahn School of Medicine at Mount Sinai, New York, NY, USA

5. Division of Infectious Diseases, Department of Medicine, Icahn School of Medicine at Mount Sinai, New York, NY, USA

6. Precision Immunology Institute, Icahn School of Medicine at Mount Sinai, New York, NY, USA

*Address:* 1 Gustave Levy Place, Annenberg, 17^th^ floor - New York, NY 10029 *Phone number:* 212-241-7393

**Figure S1.**
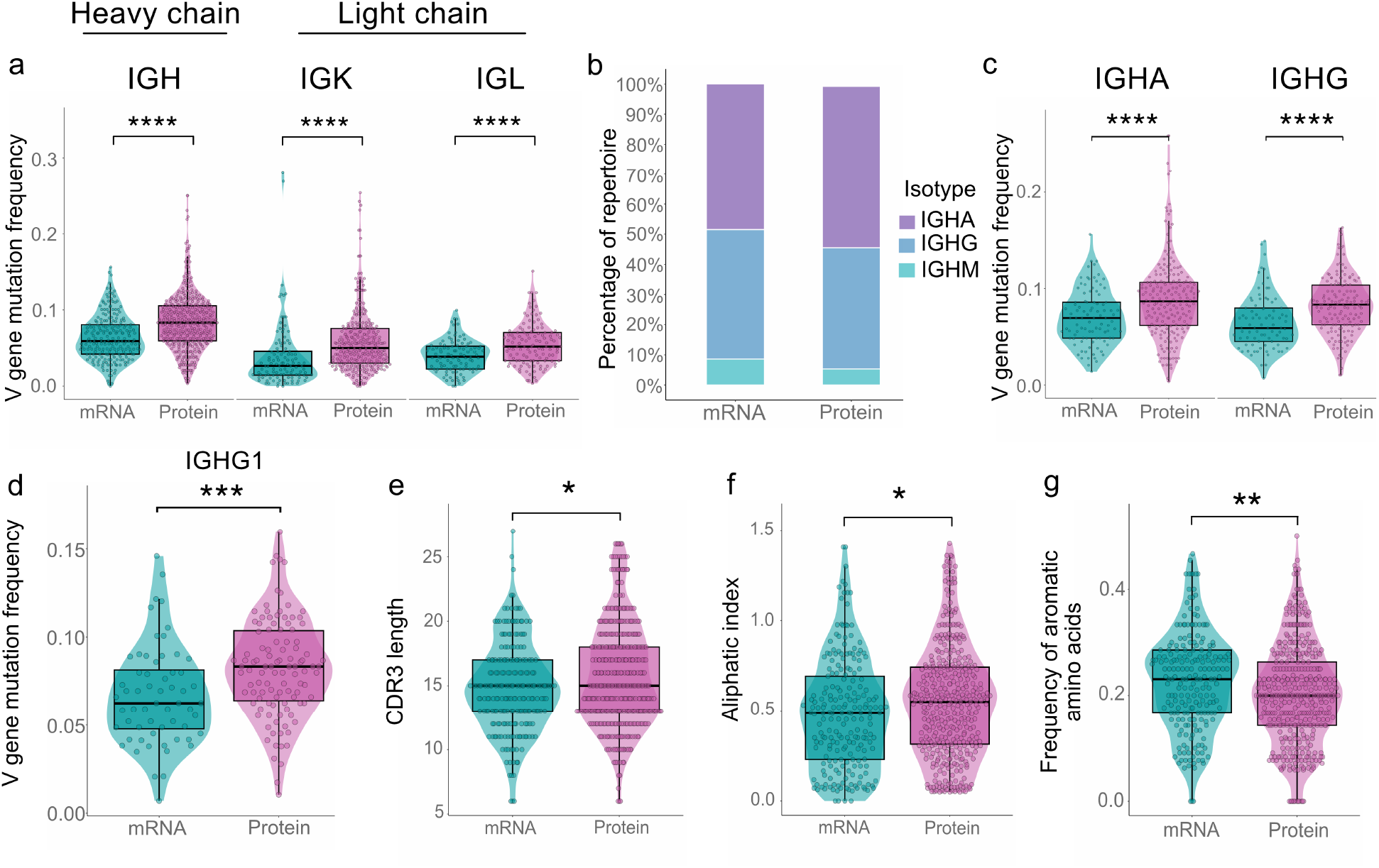
Antibody gene repertoire of mRNA- and Protein-based XBB.1.5 monovalent vaccine. (a) Frequency of nucleotide mutations in the V gene of heavy and light chain loci of plasmablasts from participants immunized with mRNA- or recombinant protein monovalent vaccines. **(b)** Antibody isotype frequency. IGHD and IGHE cells were removed from visualization. **(c)** V gene mutations in IgA and IgG sequences. **(d)** V gene mutations in IgG1. **(e)** HCDR3 length. **(f)** Relative volumes of aliphatic side chains (Ala, Val, Ile, and Leu) in the CDR3 region. **(g)** Frequency of aromatic amino acids (His, Phe, Trp, and Tyr) in the CDR3 region. Significant p-values are shown as follows: p ⩽ 0.05: *; p ⩽0.01: **; p ⩽ 0.001: ***; p ⩽ 0.0001: ****.

**Figure S2.**
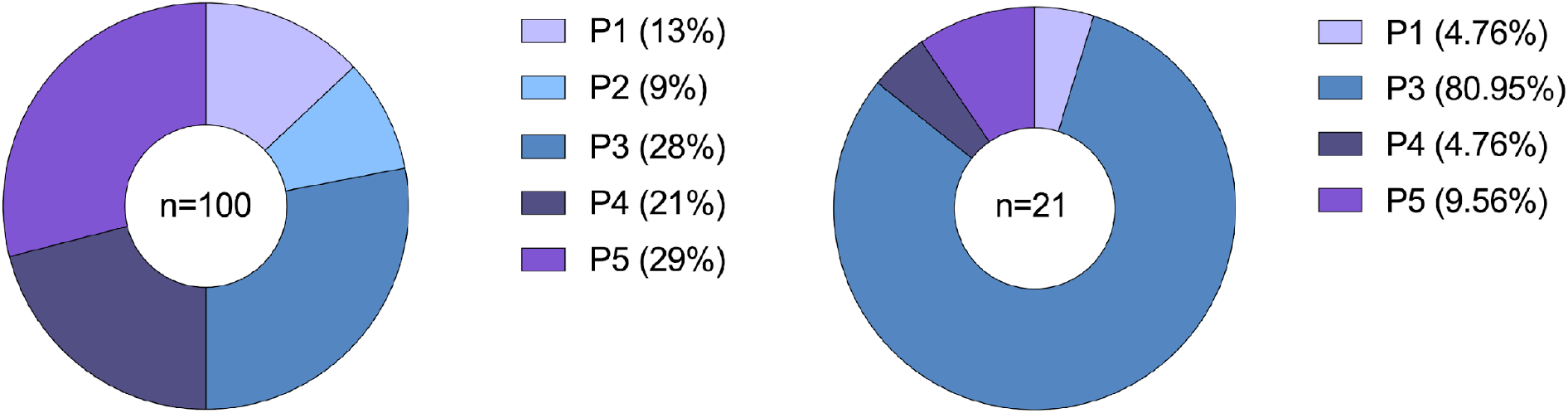
Distribution of 100 mAbs expressed across the five participants. (a) Total mAbs screened in the study. **(b)** mAbs showing binding positivity in ELISA.

**Figure S3:**
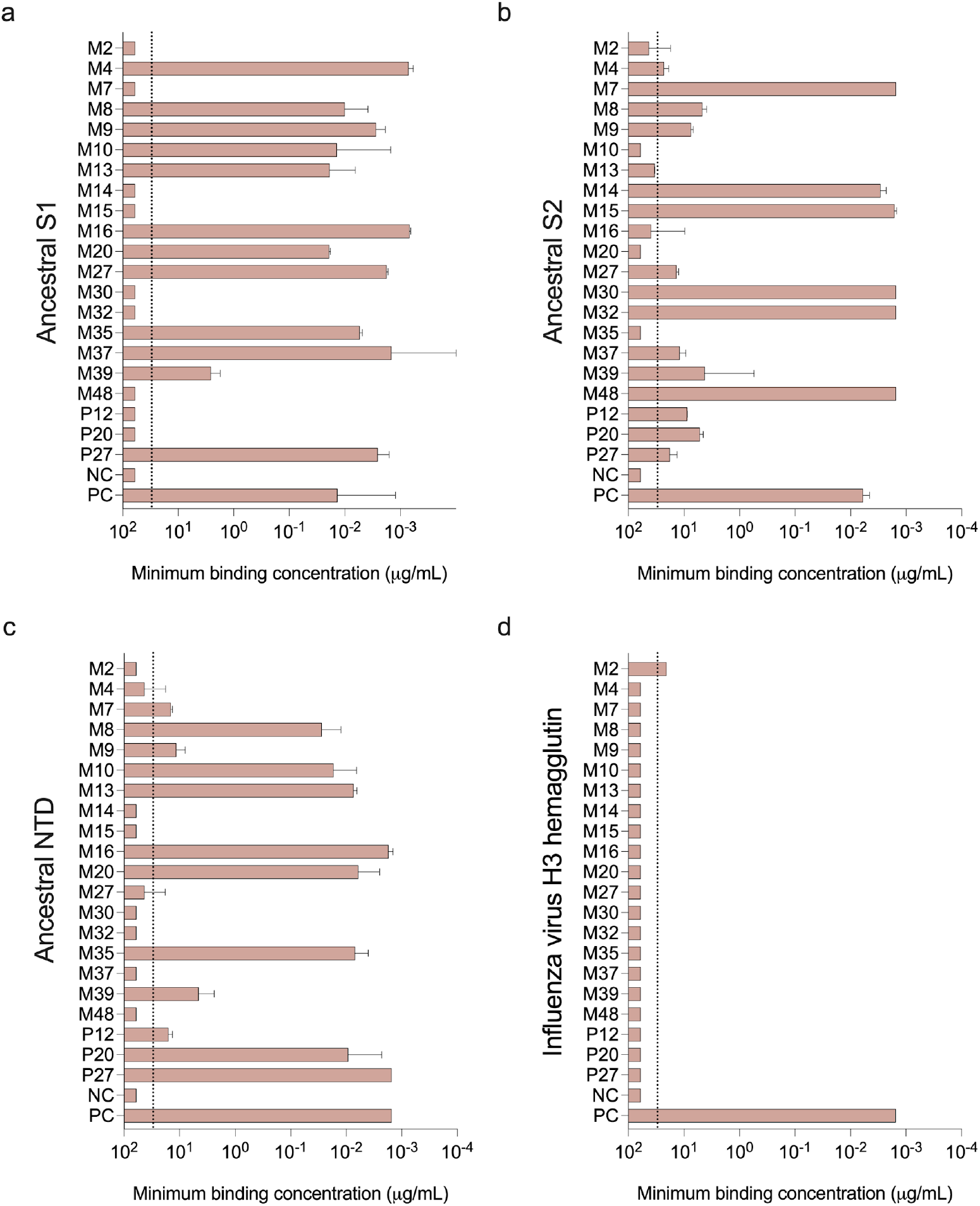
Binding capacity of monoclonal antibodies isolated from participants vaccinated with either mRNA or protein-based SARS-CoV-2 XBB.1.5 vaccines against (a) Ancestral S1, **(b)** ancestral S2, **(c)** ancestral NTD, and **(d)** an irrelevant influenza virus H3 hemagglutinin (negative control). mAbs that did not bind to any antigen were excluded from the graphs. Dashed line represents the limit of detection (LOD) which is set at the starting dilution of 30 ug/mL. Binding is defined by minimum binding concentration (µg/mL). Negative antibodies were assigned half of the LOD for graphing purposes.

**Figure S4:**
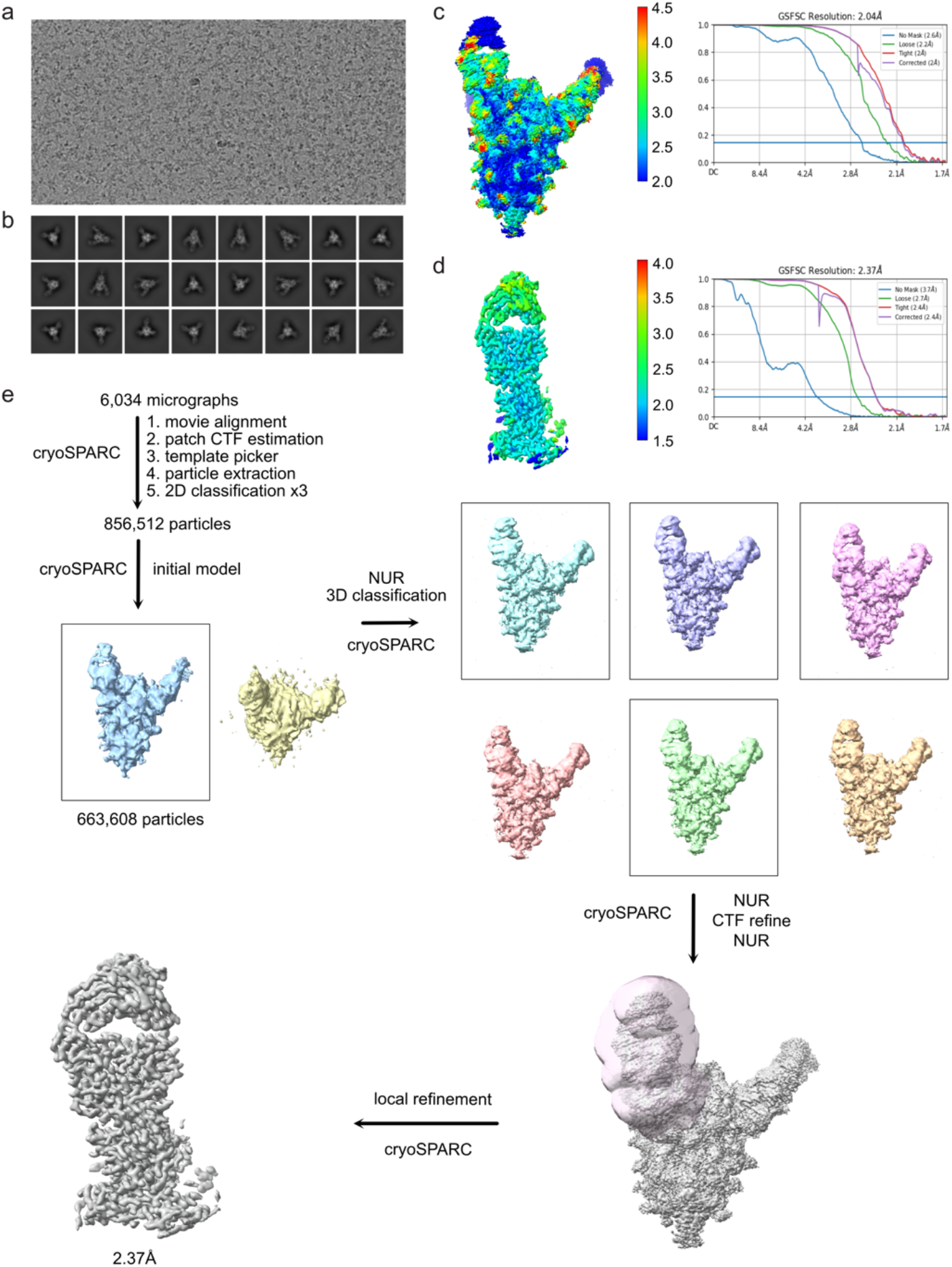
Cryo EM data processing and local resolution estimation. (a) Representative electron micrograph (**b)** 2D class average obtained for SARS-CoV-2 XBB.1.5 spike ectodomain in complex with Fab M2. Local resolution and gold-standard Fourier shell correlation curves calculated with cryoSPARC v4.3.1. **(c)** for overall (**d)** and locally refined, NTD-Fab complex. (**e)** Detailed cryo-EM data processing workflow.

**Figure S5:**
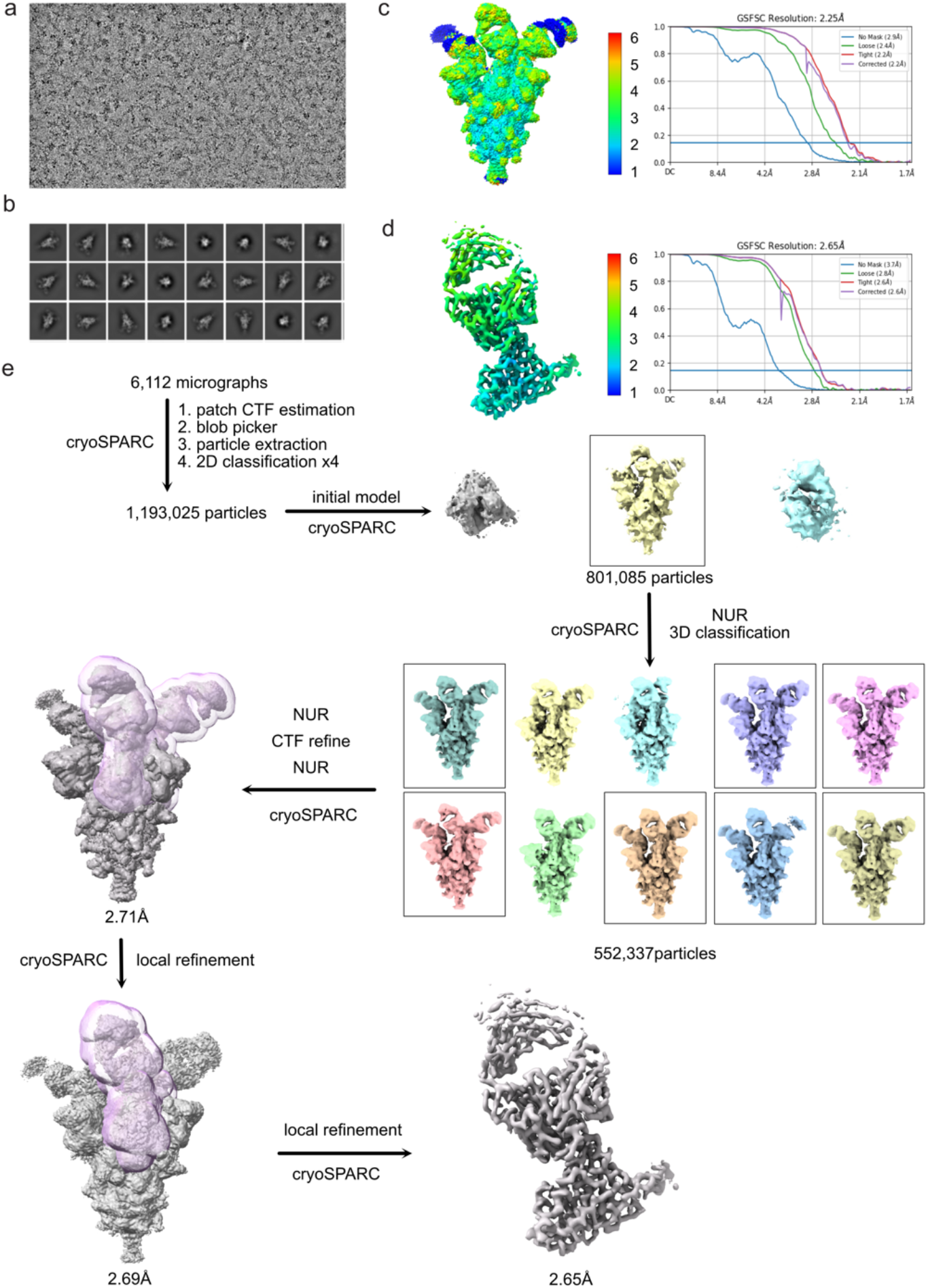
Cryo EM data processing and local resolution estimation (a) Representative electron micrograph (**b)** 2D class average obtained for SARS-CoV-2 XBB.1.5 spike ectodomain in complex with M39 Fab. Local resolution and gold-standard Fourier shell correlation curves calculated with cryoSPARC v4.3.1. **(c)** for overall (**d)** and locally refined, RBD-Fab complex. **(e)**Detailed cryo-EM data processing workflow.

**Figure S6:**
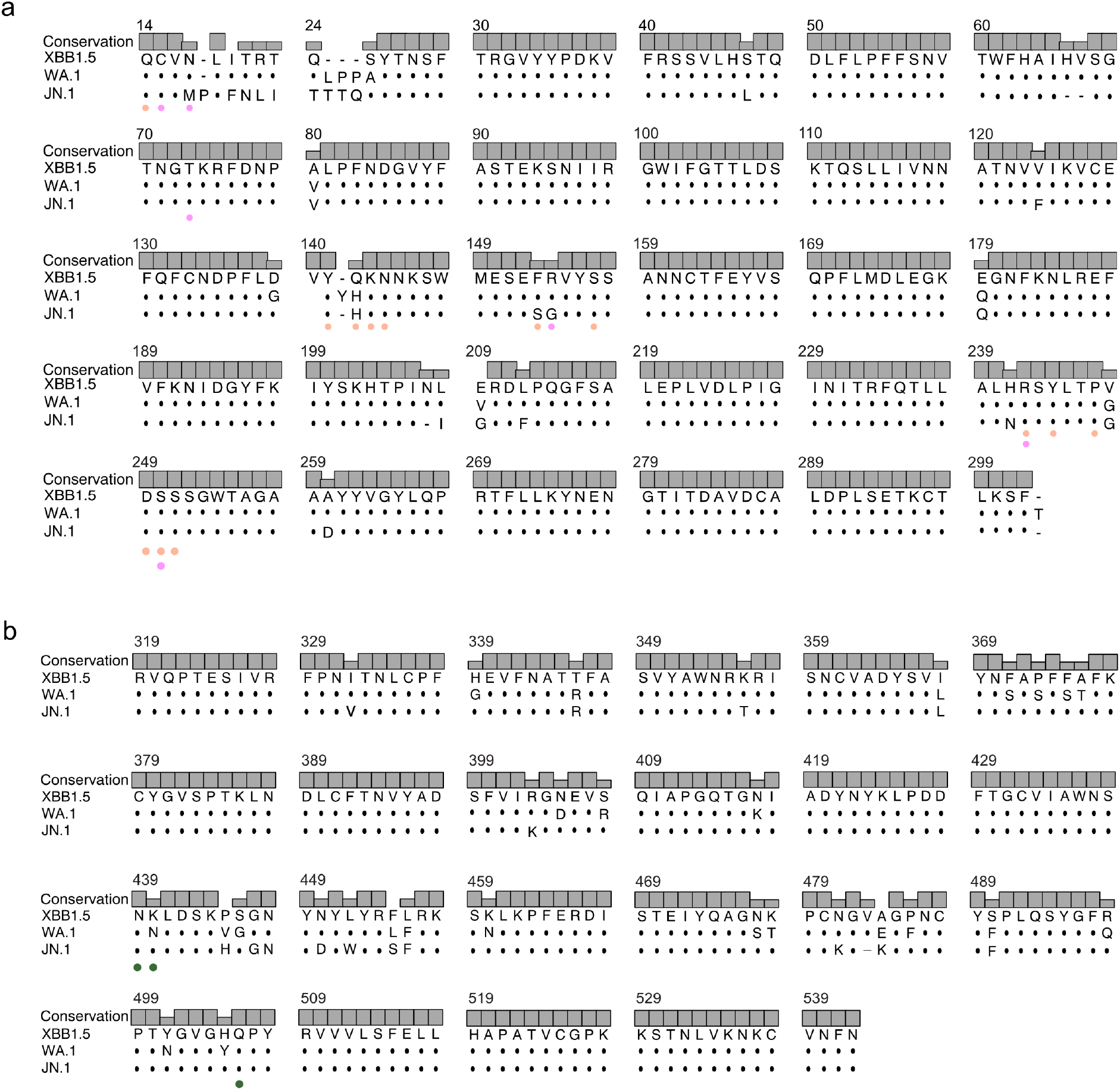
Multiple Sequence alignment **(a)** SARS-CoV-2 spike NTD (aa 14-302) of XBB.1.5, WA.1 and JN.1 **(b)** SARS-CoV-2 spike RBD (aa 319-542) of XBB.1.5, WA.1 and JN.1. Epitope residues of Fab M2 and M39 are indicated with colored dots.

**Figure S7.**
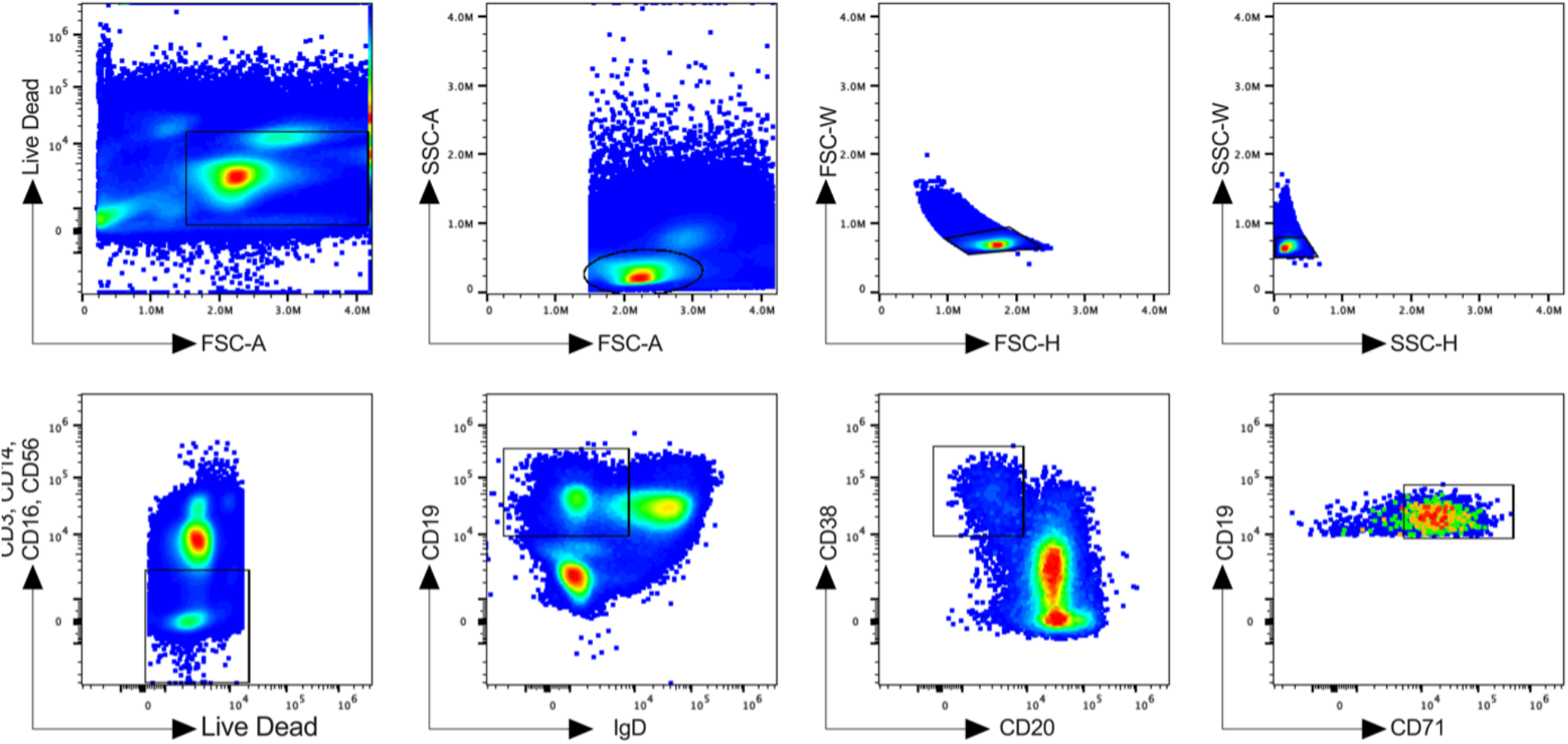
Gating strategy for plasmablasts from participant P2, 6 days post XBB.1.5 booster.

**Figure S8:**
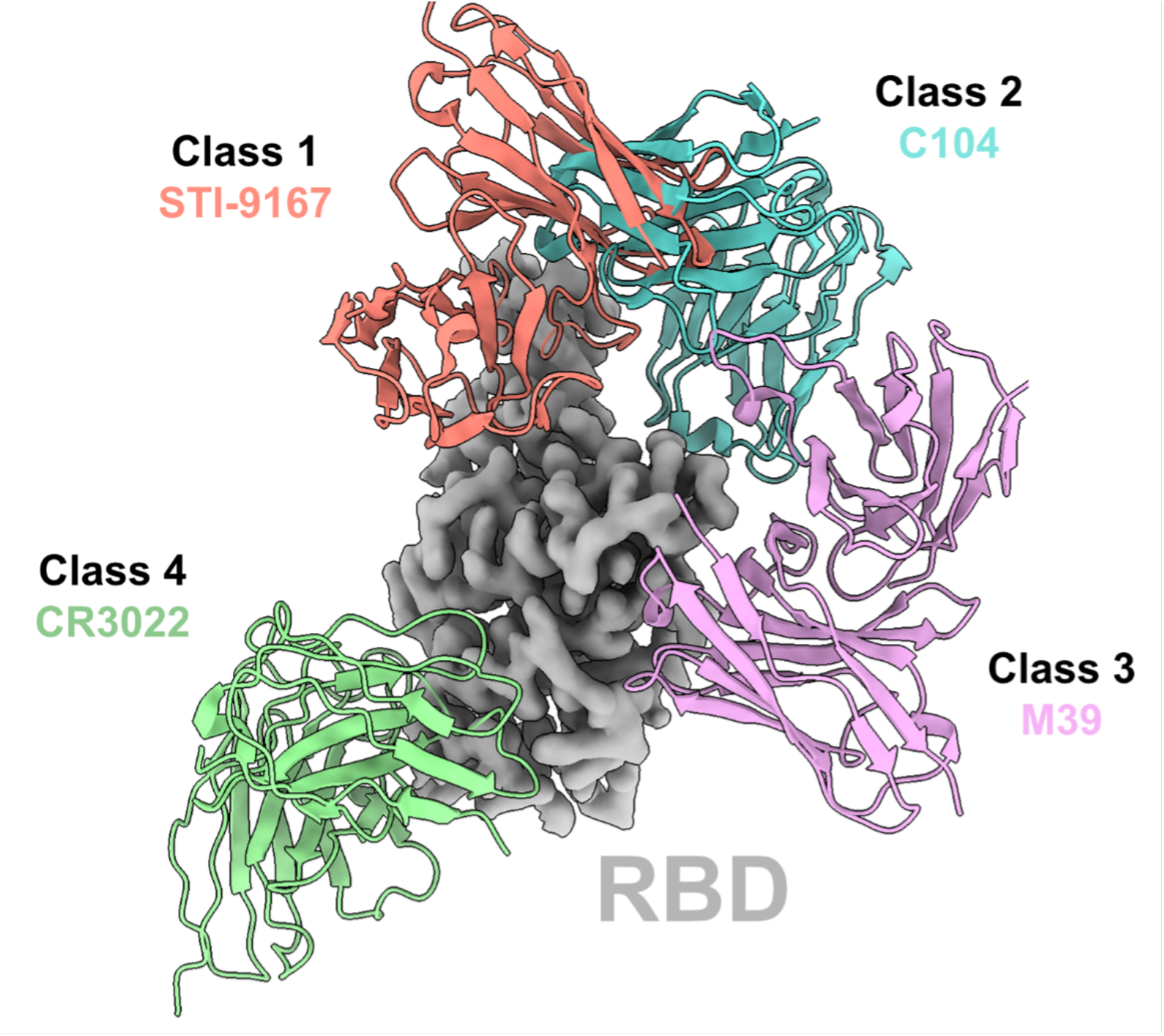
Structural illustration of representative Fabs from each class binding to its RBD epitope showing M39 as a class 3 antibody. Class 1 (STI-9167 (*62*); PDB ID 8EQF), Class 2 (C104 (*45*); PDB ID 7K8U), Class 3 (M39, PDB ID 9CCJ), Class 4 (CR3022 (*63*); PDB ID 6YLA).

## Appendix Tables

**Table S1:** Number of heavy and light chain pairs recovered from each participant.

| Participant ID | Platform | Timepoint | Number of heavy chain sequences before quality control | Number of light chain sequences before quality control | Number of heavy/light paired sequences after quality control |
| --- | --- | --- | --- | --- | --- |
| P1 | mRNA | 7 | 65 | 80 | 61 |
| P2 | mRNA | 6 | 69 | 87 | 57 |
| P3 | mRNA | 7 | 122 | 168 | 95 |
| P4 | Protein | 7 | 205 | 220 | 201 |
| P5 | Protein | 7 | 194 | 220 | 189 |

**Table S2:** Viability (in %) of human PBMCs used in sorting experiments.

| Participant ID | Days post-vax | Cell type | Viability |
| --- | --- | --- | --- |
| P1 | 7 | Plasmablasts | 94% |
| P2 | 6 | Plasmablasts | 98% |
| P3 | 7 | Plasmablasts | 98% |
| P4 | 7 | Plasmablasts | 97% |
| P5 | 7 | Plasmablasts | 97% |

**Table S3:** Antibodies Used for Flow Cytometry.

| Antibody | Vendor | Catalog Number | Dilution |
| --- | --- | --- | --- |
| CD19 BUV563 | BD Biosciences | 612916 | 1:200 |
| IgD Pacific Blue | Biolegend | 348224 | 1:50 |
| CD20 BV510 | Biolegend | 302340 | 1:100 |
| CD3 PerCP | Biolegend | 344814 | 1:100 |
| CD14 PerCP | Biolegend | 367152 | 1:200 |
| CD16 PerCP | Biolegend | 302030 | 1:200 |
| CD56 PerCP | Biolegend | 318342 | 1:200 |
| CD38 AF810 | Biolegend | 334120 | 1:200 |
| CD71 PE/Dazzle594 | Biolegend | 334120 | 1:200 |

**Table S4:** Cryo-EM data collection and model validation statistics.

| Data Collection |  |  |
| --- | --- | --- |
|  | PDB ID 9CCJ and EMD-45444 | PDB ID 9CCI and EMD-45443 |
| Grid type | UltrAuFoil gold R1.2/1.3 | UltrAuFoil gold R1.2/1.3 |
| Microscope/voltage/detector | Titan Krios/300 kV/Gatan K3 summit | Titan Krios/300 kV/Gatan K3 summit |
| Magnification | 105,000 | 105,000 |
| Recording mode | counting | counting |
| Total dose | 49.883 e-/Å<sup>2</sup>/s | 49.883 e-/Å<sup>2</sup>/s |
| Pixel size | 0.825 Å/pixel | 0.825 Å/pixel |
| Defocus range | 0.6 to 2.4 µm | 0.6 to 2.5 µm |
| No. micrographs used | 6,034 | 6,112 |
| Total particles picked | 856,516 | 1,193,025 |
| Model Validation |  |  |
| Composition (#) |  |  |
| Chains | 3 | 3 |
| Atoms | 5410 | 4921 |
| Residues | Protein: 686 Nucleotide: 0 | Protein: 631 Nucleotide: 0 |
| Ligands | NAG: 7, BMA:1 | NAG: 1 |
| Bonds (RMSD) |  |  |
| Length (Å) (# > 4sigma) | 0.008 (2) | 0.005 (0) |
| Angles (°) (# > 4sigma) | 1.346 (11) | 0.871 (2) |
| MolProbity score | 1.40 | 1.21 |
| Clash score | 2.35 | 1.66 |
| Ramachandran plot (%) |  |  |
| Outliers | 0.30 | 0.00 |
| Allowed | 5.38 | 4.19 |
| Favored | 94.32 | 95.81 |
| Rotamer outliers (%) | 0.00 | 0.00 |
| Cbeta outliers (%) | NA | NA |
| Peptide plane (%) |  |  |
| Cis proline/general | 7.3/0.0 | 5.3/0.0 |
| Twisted proline/general | 0.0/0.0 | 0.0/0.0 |
| CaBLAM outliers (%) | 2.75 | 2.29 |
| Data |  |  |
| Lengths (Å) | 74.25, 93.22, 127.05 | 72.60, 97.35, 142.72 |
| Angles (°) | 90.00, 90.00, 90.00 | 90.00, 90.00, 90.00 |
| Supplied Resolution (Å) | 2.4 | 2.6 |
| Resolution Estimates (Å) | Masked | Masked |
| d FSC (half maps; 0.143) | 2.4 | 2.6 |
| d 99 (full/half1/half2) | 2.0/1.7/1.7 | 2.2/1.9/1.9 |
| d model | 1.6 | 1.7 |
| **D FSC model (0/0.143/0.5)** | 1.4/1.5/2.2 | 1.4/1.6/3.0 |
| **Map min/max/mean** | -0.00/1.83/0.01 | -0.00/1.66/0.02 |
| **Model vs. Data** |  |  |
| **CC (mask)** | 0.84 | 0.81 |
| **CC (box)** | 0.81 | 0.58 |
| **CC (peaks)** | 0.78 | 0.39 |
| **CC (volume)** | 0.84 | 0.81 |
| **Mean CC for ligands** | 0.67 | 0.69 |

